# Pupil dilation and microsaccades provide complementary insights into the dynamics of arousal and instantaneous attention during effortful listening

**DOI:** 10.1101/2023.02.06.527294

**Authors:** Claudia Contadini-Wright, Kaho Magami, Nishchay Mehta, Maria Chait

**Affiliations:** Ear Institute, University College London, London, UK; UCLH Biomedical Research Centre, Hearing Health theme; Royal National ENT and Eastman Dental Hospitals, UCLH

## Abstract

Listening in noisy environments requires effort – the active engagement of attention and other cognitive abilities, as well as increased arousal. The ability to separately quantify the contribution of these components is key to understanding the dynamics of effort and how it may change across listening situations and in certain populations. We concurrently measured two types of ocular data in young participants (both sexes) - Pupil dilation (PD) (thought to index arousal aspects of effort) and Microsaccades (MS) (hypothesized to reflect automatic visual exploratory sampling) whilst listeners were performing a speech-in-noise task under high- (HL) and low-(LL) perceptual load conditions. Sentences were manipulated so that the behaviourally relevant information (keywords) appeared at the end (Exp1) or beginning (Exp2) of the sentence, resulting in different temporal demands on focused attention. In line with previous reports, PD effects were associated with increased dilation under load. We observed a sustained difference between HL and LL conditions, consistent with increased phasic and tonic arousal. Importantly we show that MS rate was also modulated by perceptual load, manifested as a reduced MS rate in HL relative to LL. Critically, in contrast to the sustained difference seen for PD, MS effects were localised in time, specifically during periods when demands on auditory attention were greatest. These results demonstrate that auditory selective attention interfaces with the mechanisms controlling MS-generation, establishing MS as an informative measure, complementary to PD, with which to quantify the temporal dynamics of auditory attentional processing under effortful listening conditions.

**Significance Statement:** Listening effort, reflecting the “cognitive bandwidth” deployed to effectively process sound in adverse environments, contributes critically to listening success. Understanding listening effort and the processes involved in its allocation is a major challenge in auditory neuroscience. Here we demonstrate that the microsaccade rate can be used to index a specific sub-component of listening effort - the allocation of instantaneous auditory attention - that is distinct from the modulation of arousal indexed by pupil dilation (currently the dominant measure of listening effot). These results reveal the push-pull process through which auditory attention interfaces with the (visual) attention network that controls microsaccades, establishing microsaccades as a powerful tool for measuring auditory attention and its deficits.

## Introduction

Listening in noisy environments draws on cognitive capacities, such as attention and memory, that support the tracking of relevant signals among interference (Peelle, 2018). The active allocation of these resources is termed “listening effort” (Pichora-Fuller, 2016). Better insight into listening effort and its neural underpinnings is critical for understanding the challenges faced by listeners under adverse conditions, and for managing the breakdown of these processes due to aging, hearing loss, or certain neurological conditions.

Effortful listening to speech in noise engages heightened arousal as well as increased demands on attention, working memory, and linguistic processing (Peelle, 2018). Unravelling these various aspects - in particular separating arousal-related effects from those related to the allocation of cognitive resources - is a key challenge for the field (Zekveld et al, 2014; White & Langdon, 2021; Ritz et al, 2021; Haro et al, 2022; McGarrigle et al, 2021). Correlations between subjective, behavioral, and physiological measures of listening effort have not always yielded consistent results (Alhanbali et al., 2019), implying that different approaches may be capturing distinct facets of this construct.

Pupillometry is frequently used to measure the arousal component of listening effort (Zekveld et al, 2018; Ohlenforst et al, 2018; Winn et al, 2018). Non-luminance mediated pupil dilation (PD) is linked to activity in the Locus Coeruleus (LC), which supplies Norepinephrine to the central nervous system and therefore controls vigilance and arousal (Wang & Munoz, 2015; Joshi et al, 2016). Stimulus-evoked PD is related to *phasic* activity within the LC, associated with instantaneous arousal, while baseline changes in pupil size are hypothesized to index *tonic* LC activity, associated with sustained alertness and engagement. Previous literature has consistently demonstrated that conditions associated with greater listening effort often lead to an increase in pupil dilation (relative to baseline) (Zekveld et al 2018; Winn et al, 2018). These effects are commonly extended in time, with peak PD frequently occurring after sentence offset, hypothesized to reflect a ‘release’ of arousal at the conclusion of the perceptual process (Winn et al., 2016; Winn & Moore, 2018; Winn et al., 2022). Conditions of high listening load have also been associated with increased *baseline* pupil size, proposed to reflect a balance between sustained arousal and build-up of mental fatigue (Hopstaken, et al, 2015; McGarrigle et al, 2017).

Here, we hypothesize that a specific component of listening effort, the allocation of instantaneous attention, which is distinct from the state of arousal reflected by PD, can be indexed by measuring another type of ocular activity -microsaccades.

Microsaccades (MS) are small fixational eye movements controlled by a network encompassing the frontal eye fields and the superior colliculus (Hafed et al, 2015; Rucci & Victor, 2015) and are hypothesized to reflect unconscious continuous exploration of the visual environment. Recent findings suggest that this sampling is affected by the attentional state of the individual: MS incidence reduces during -and in anticipation of-task-relevant events (Widmann et al, 2014; Denison et al, 2019; Abeles et al, 2020) and under high load (Dalmaso et al, 2017; Yablonski et al, 2017; Lange et al, 2017). Together, this evidence suggests that the processes generating MS draw on shared, limited computational capacity such that MS-indexed visual exploration is reduced when attentional resources are depleted by other perceptual tasks. Despite the richness of information potentially conveyed by MS, our understanding of how auditory perceptual processes might interface with the attentional sampling mechanisms that control MS is limited.

We recorded PD and MS concurrently while participants listened to sentences in noise under conditions of high and low listening effort (Figure 1). The placement of relevant keywords was manipulated to control the timing of instantaneous attentional engagement. We expected both PD and MS to be modulated by load, but exhibiting a different timing profile: temporally extended effects of PD-linked arousal, but transient and time-specific effects of MS incidence - precisely during periods (when keywords are presented) where the demands on auditory attention are highest.

**Figure 1:**
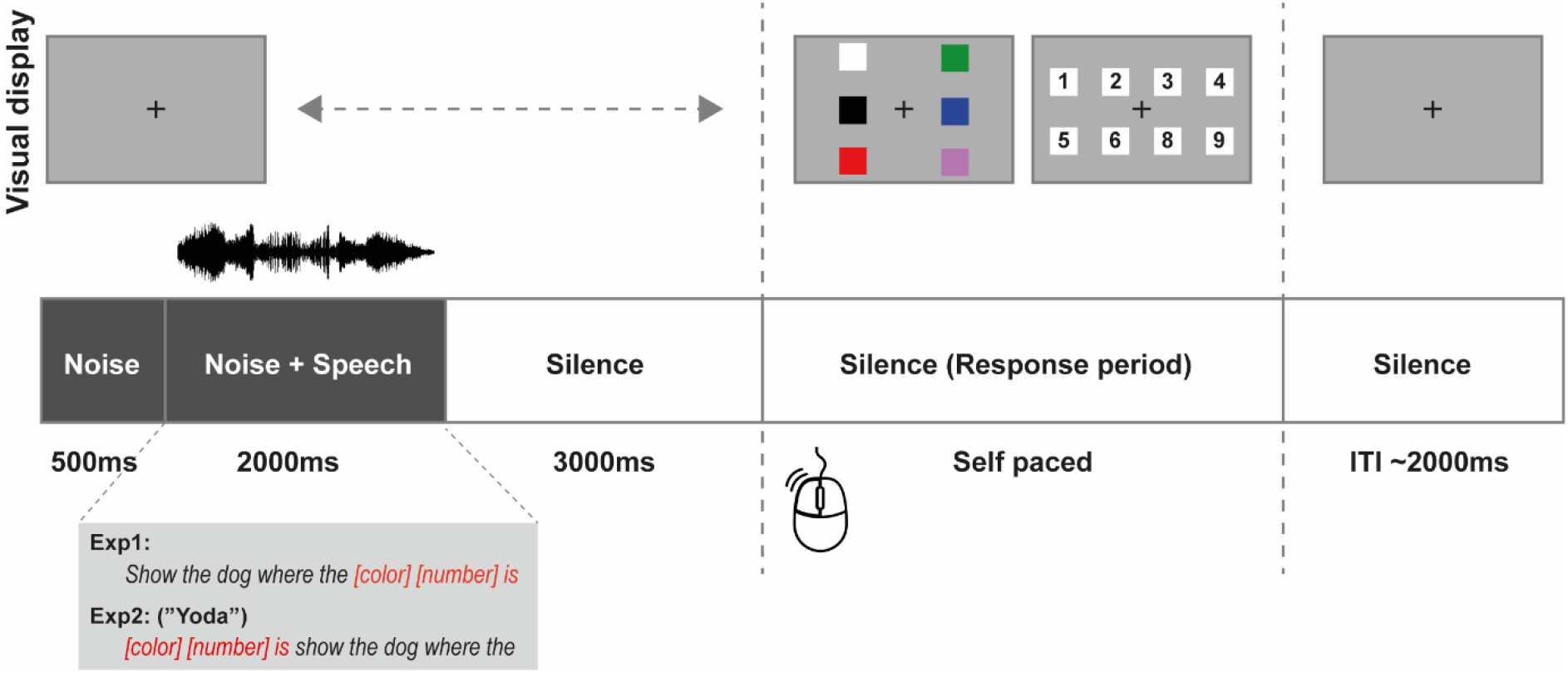
Trial structure. In Exp1, sentences were in the form “show the dog where the [color] [number] is”. In Exp2 (‘Yoda’), sentence structure was reversed such that the keywords ([color] [number]) appeared first. Trials began with 0.5s of noise only, followed by the onset of the sentence (∼2s) and then a silent period (3s). A response display then appeared on the screen. Participants logged their responses by selecting the correct color, then number. Visual feedback was provided. The simple sentential material and response procedure reduced demands on working memory and executive function, and emphasized the draw on attentional resources required to “fish out” the keywords from the noisy signal.

## Methods

### Participants

All experimental procedures were approved by the research ethics committee of University College London, and written informed consent was obtained from each participant.

All participants were native or long-term fluent English speakers. All reported normal hearing with no history of otological or neurological disorders. Participants reported normal or corrected-to-normal vision, with Sphere (SPH) prescriptions no greater than 3.5. 35 participants were recruited and underwent testing in both experimental designs. 2 different sets of participants were recruited for the 2 experiments.

**Exp1:** 35 paid participants between the ages of 18-35 took part (30 female, mean age= 23.49, SD = 4.09). Four participants were eventually excluded from the dataset, 2 due to substantial amounts of missing pupil data (blinks and/or gazes away from fixation; see ***Pupillometry Preprocessing and Analysis***) and 2 due to poor behavioral performance, resulting in a final set of 31 participants (27 female, mean age= 23.45, SD = 3.87).

**Exp2:** 35 paid participants between the ages of 18-35 took part (28 female, mean age = 25.26, SD = 5.24). Four participants were eventually excluded from the dataset, 3 due to substantial amounts of missing pupil data and 1 due to poor behavioral performance, resulting in a final set of 31 participants (25 female, mean age = 25.45, SD= 4.97).

### Design and Materials

The experimental session lasted approximately 1.5 hours and was comprised of three stages:

#### (1) Threshold estimation

A speech-in-noise reception threshold was first obtained from each participant, using the CRM task (see **Threshold estimation** subsection below). We used an adaptive procedure to determine the 50% correct threshold.

#### (2) Baseline pupil measures

Prior to the main experimental session, we performed a series of brief baseline measures of pupil reactivity (Light reflex, Dark reflex etc). These included measuring pupil responses to a slow, gradual change in screen brightness; to a sudden flashing white screen; to a sudden flashing black screen; and to a sudden presentation of a brief auditory stimulus (harmonic tone). These measurements were used to confirm normal pupil responsivity and to identify outlying participants (none here).

#### (3) Main experiment

In the main experiment participants performed two blocks of the CRM task while their ocular data were being recorded. In one of the blocks (‘High load’) the signal-to-noise ratio (SNR) was set to the threshold obtained in (1), simulating a difficult listening environment. In the second block (‘Low load’) the SNR was set to the threshold obtained in (1) plus 10 dB to create a much easier listening environment (as in McGarrigle et al, 2020). The order of the two blocks was counterbalanced across participants.

All experimental tasks were implemented in MATLAB and presented via Psychophysics Toolbox Version 3 (PTB-3).

##### Threshold estimation

Auditory stimuli were sentences introduced by Messaoud-Galusi, Hazan, & Rosen (2011) –which are a modified version of the CRM (“Coordinate Response Measure”) corpus described by Bolia et al., 2000. Sentences in Experiment 1 (including threshold estimation) were in the form “Show the dog where the [color] [number] is”. Sentences in Experiment 2 (including threshold estimation) were in the form “[color] [number] is show the dog where the”. The colors that could appear within a target sentence were black, red, white, blue, green and pink. The numbers could be any digit from 1-9 with the exception of 7, as its bisyllabic phonetic structure makes it easier to identify. Consequently, there were a total of 48 possible combinations of color and number. Sentence duration ranged between 1.9 and 2.4s, with the majority having a duration of 2.1s. Sentences were embedded in Gaussian noise. The overall loudness of the noise+speech mixture was fixed at ∼70dB SPL. The SNR between the noise and speech was initially set to 20dB, and was adjusted using a one-up-one-down adaptive procedure, tracking the 50% correct threshold. Initial steps were of 12dB SNR, and decreased steadily following each reversal up to a minimum step size of 2dB. The test ended after 7 reversals or after a total of 25 trials and took about 2 minutes to complete. The speech reception threshold was calculated as the mean SNR of the final four reversals. Participants completed ∼3 runs in total (the first was used as a practice). The threshold obtained from the final run was used for the ‘High load’ condition in the main experiment. The threshold plus 10dB was used for the ‘Low load’ condition in the main experiment.

##### Main task

In the main experiment (‘High load’ (HL) and ‘Low load’ (LL) blocks; 15 min total), the same stimuli were used as for threshold estimation, but the SNR was fixed as described above. Each block contained 30 trials. Participants fixated on a black cross presented at the centre of the screen (grey background). The structure of each trial is schematized in Figure 1. Trials began with 0.5s of noise, followed by the onset of the sentence in noise (∼2s long) and then a silent period (3s). A response display then appeared on the screen and participants logged their responses to the task by selecting the correct color first, then the number, using a mouse. Visual feedback was provided. At the end of each trial, participants were instructed to re-fixate on the cross in anticipation of the next stimulus.

##### Procedure

Participants sat with their head fixed on a chinrest in front of a monitor (24-inch BENQ XL2420T with a resolution of 1920×1080 pixels and a refresh rate of 60 Hz) in a dimly lit and acoustically shielded room (IAC triple walled sound-attenuating booth). They were instructed to continuously fixate on a black cross presented at the center of the screen against a grey background. An infrared eye-tracking camera (Eyelink 1000 Desktop Mount, SR Research Ltd.) placed below the monitor at a horizontal distance of 62cm from the participant was used to record pupil data. Auditory stimuli were delivered diotically through a Roland Tri-capture 24-bit 96 kHz soundcard connected to a pair of loudspeakers (Inspire T10 Multimedia Speakers, Creative Labs Inc, California) positioned to the left and right of the eye tracking camera. The loudness of the auditory stimuli was adjusted to a comfortable listening level for each participant. The standard five-point calibration procedure for the Eyelink system was conducted prior to each experimental block and participants were instructed to avoid any head movement after calibration. During the experiment, the eye-tracker continuously tracked gaze position and recorded pupil diameter, focusing binocularly at a sampling rate of 1000 Hz. Participants were instructed to blink naturally during the experiment and encouraged to rest their eyes briefly during inter-trial intervals. Prior to each trial, the eye-tracker automatically checked that the participants’ eyes were open and fixated appropriately; trials would not start unless this was confirmed.

### Pupillometry Preprocessing and Analysis

Only data from the left eye was analyzed. Intervals when the participant gazed away from fixation (outside of a radius of 100 pixels around the center of the fixation cross) or when full or partial eye closure was detected (e.g. during blinks) were automatically treated as missing data. Participants with excessive missing data (>50%) were excluded from further analysis. This applied to two of the participants in Experiment 1 and three participants in Experiment 2.

#### PD to speech

The pupil data from each trial were epoched from – 2s to 5.5 sec from sound (noise) onset. Epochs with more than 50% missing data were discarded from the analysis. On average, <1 trial was discarded per subject in each of the experimental blocks. Missing data in the remaining trials were recovered using shape-preserving piecewise linear interpolation. Trials where 10% or more of the data were identified as outlying (>3SD from the condition mean) were removed from analysis. On average, <1 trial was discarded per subject in each of the experimental blocks. Data were then z-scored (across all trials, collapsed across the High and Low load conditions) for each participant, and time-domain-averaged across all epochs of each condition to produce a single time series per condition. Both baseline corrected (from 0.2-0s pre onset) and non-baseline corrected data are reported.

#### Microsaccade Preprocessing and Analysis

Microsaccade (MS) detection was based on an approach proposed by Engbert & Kliegl (2003). MS were extracted from the continuous eye-movement data based on the following criteria: (1) A velocity threshold of λ = 6 times the median-based standard deviation within each condition (2) Above-threshold velocity lasting between 5ms and 100ms (3) The events are detected in both eyes with onset disparity <10ms (4) The interval between successive microsaccades is longer than 50ms. Extracted microsaccade events were represented as unit pulses (Dirac delta) and epoched as described for the PD analysis above. In Experiment 1, the eye-tracker settings (set to briefly interrupt the recording prior to the onset of each trial) resulted in a short period (several samples) of lost data 0.3 – 0s prior to stimulus onset. To address the consistent artefactual absence of MS during that interval, these data were replaced by 300ms of data between 0.6-0.3s pre-onset selected from a random trial of the same participant.

In each condition, for each participant, the event time series were summed and normalized by the number of trials and the sampling rate. Then, a causal smoothing kernel *ω*(*τ*) = *α*^2^×*τ*×*e*^−*ατ*^ was applied with a decay parameter of 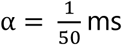 (Dayan & Abbott, 2001; Rolfs et al., 2008; Widmann et al., 2014), paralleling a similar technique for computing neural firing rates from neuronal spike trains (Dayan & Abbott, 2001; see also Joshi et al., 2016; Rolfs et al., 2008). To account for the time delay caused by the smoothing kernel, the time axis was shifted by the peak of the kernel window.

#### Statistical analysis

To identify time intervals in which a given pair of conditions exhibited PD/MS differences, a non-parametric bootstrap-based statistical analysis was used (Efron & Tibshirani, 1994). The difference time series between the conditions was computed for each participant and these time series were subjected to bootstrap re-sampling (1000 iterations; with replacement). At each time point, differences were deemed significant if the proportion of bootstrap iterations that fell above or below zero was more than 99% (i.e. p<0.01). The analysis was conducted on the full epoch as plotted and all significant time points are shown.

## Results

### Behavioral Performance

Figure 2 shows the behavioral performance on the CRM task in Experiments 1 and 2. Speech material in Experiment 1 consisted of ‘standard’ CRM sentences (“Show the dog where the [color] [number] is”). All sentences were initially identical (“Show the dog where the…..”), allowing the listener to slowly increase arousal and attention as they prepared to identify the keywords, which always occurred at the end. In contrast, Experiment 2 used the same stimuli, but with the keywords moved to sentence onset (“[color] [number] is show the dog where the”). We refer to this condition as “Yoda” because it is similar to the speech pattern of the iconic Star Wars character (LaFrance, 2015). This material required rapid focusing of attention immediately at sentence onset, but allowed for potential disengagement of attention and arousal partway through the sentence as the remaining speech was not relevant for the task. Additionally, the unusual grammatical structure required adjusting to. Despite these distinctions, we did not observe significant differences in behavioral performance between experiments.

**Figure 2:**
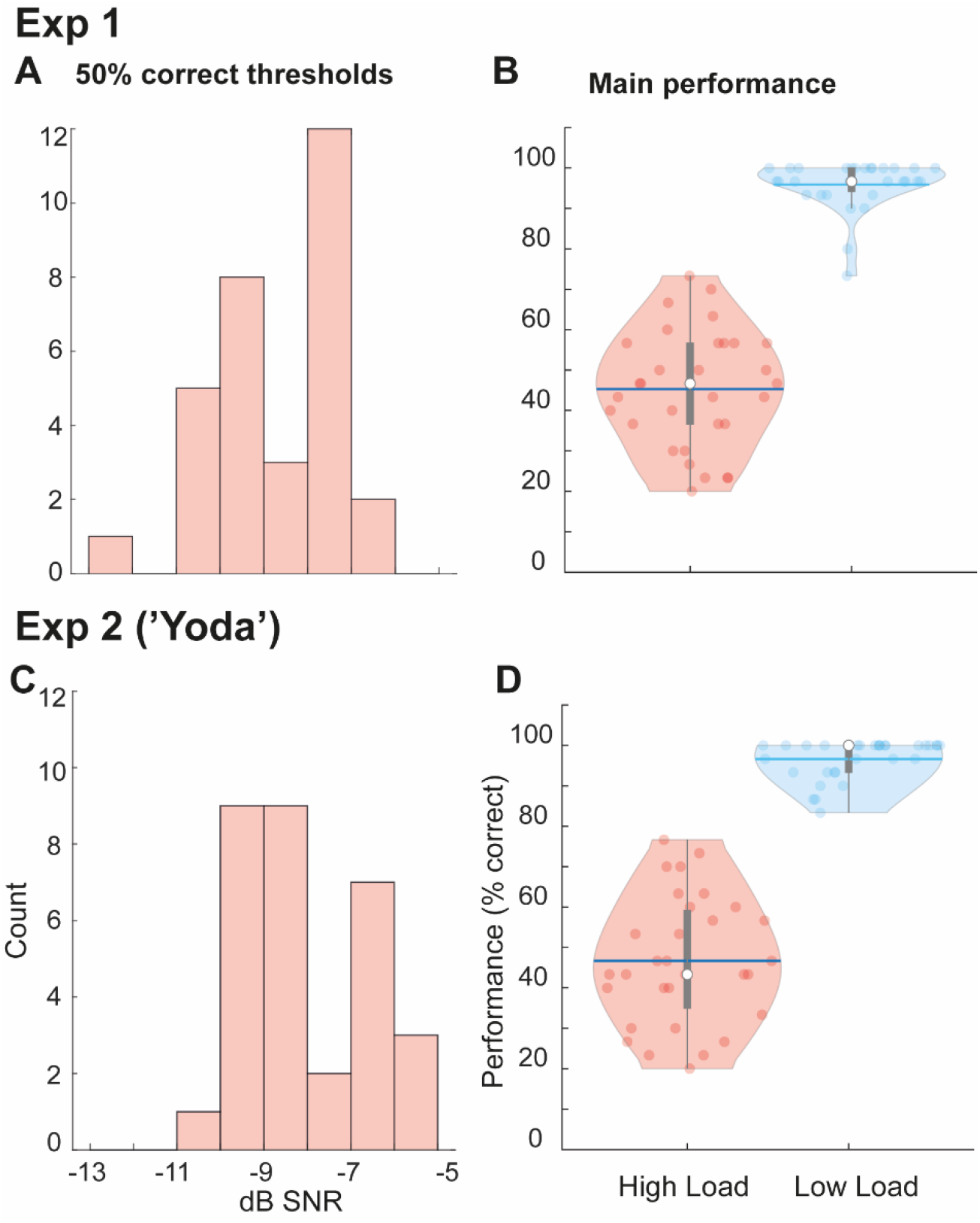
Speech-in-noise performance in Experiments 1 and 2. [A,C] a histogram of 50% correct SNRs; lower values indicated higher sensitivity to speech in noise. [B,D] main task performance in the High and Low Load blocks. The horizontal blue lines represent the mean, while the white dots represent the median.

The left panels plot histograms of the 50% correct SNRs measured for each participant - these values were used for the HL condition in the main experiment. The right panels plot the performance in the main experiment (scored in %). In the HL condition, participants achieved an average of 46.34% (Experiment 1) and 46.67% (Experiment 2) correct responses, which rose to a mean of 96.02% (Experiment 1) and 96.67% (Experiment 2) in the LL condition. This confirms that the load manipulation was successful. A Mann-Whitney U test confirmed no difference between SNRs across experiments (U=598, *p*=0.097). Similarly, an analysis of the main performance data confirmed no main effect of experiment (F(1,60)=0.225, *p*=0.64), only a main effect of load (F(1,60)=787.34, *p*<0.001).

### Experiment 1: Perceptual load modulates pupil size

Focusing on the load manipulation in Experiment 1, Figure 3 plots the mean PD to the target sentences in noise; z scored (across HL and LL conditions for each participant) and averaged across participants.

**Figure 3:**
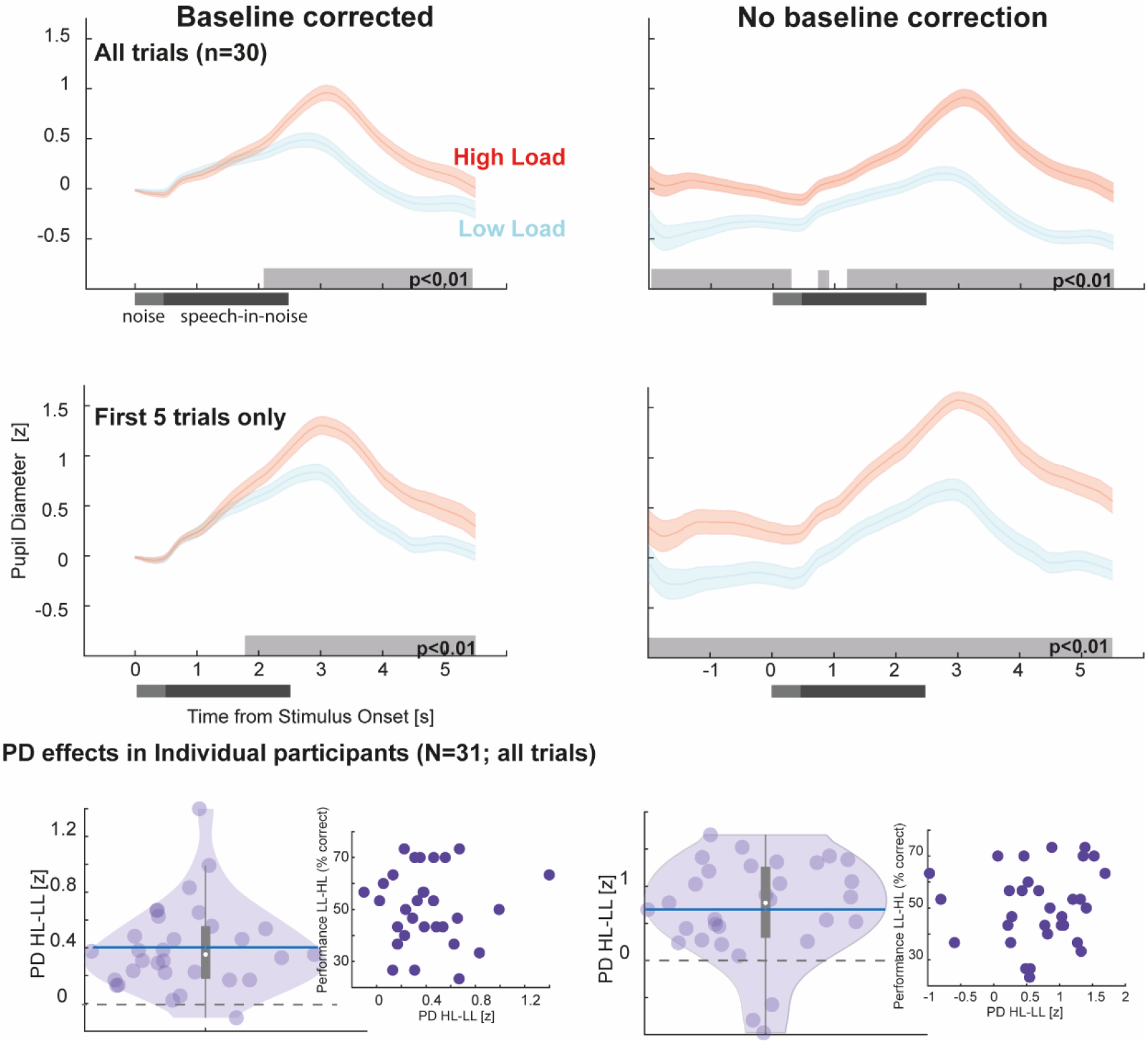
Pupil diameter is robustly modulated by listening load. [Top] Pupil diameter was consistently larger in the HL relative to the LL condition. This was also observable when analysing the first 5 trials of each condition, confirming a robust effect. Significant differences between conditions are indicated by the grey horizontal lines. [Bottom] Load effect in each participant (quantified by taking the difference between HL and LL Pupil Dilation Response (PDR) in the interval 2-5s post onset). Baseline corrected data on the left; non-baseline corrected on the right. Horizontal blue line represents the mean, while the white dot represents the median. Insets show the PD difference (horizontal axis) against performance difference between the LL and HL conditions.

To quantify the sentence evoked pupil dilation, we baseline corrected the responses (0.2s pre-noise onset; Fig 3 left). The data demonstrate a clear increase in pupil size elicited by the target sentence. In both conditions, pupil diameter rose monotonically from sentence onset, reaching its peak following sentence offset (3.09s post-onset in HL; 2.78s post-onset in LL). This is consistent with previous reports suggesting that the monotonic increase in PD reflects the gradual build-up of arousal and its subsequent release after the conclusion of perceptual processing (Winn et al., 2022; Winn & More, 2018; Winn et al., 2016). This increase was significantly larger in HL relative to LL, with the difference emerging from 2s post-onset and persisting to the end of the epoch. The non-baseline corrected responses (Fig 3; right) additionally revealed a tonic difference between HL and LL conditions that is sustained throughout the trial. This is consistent with previous observations of increased baseline pupil size under conditions of effortful listening, likely reflecting increased sustained arousal.

The effect of load on PD was observable in the majority of participants (Figure 3 bottom) and also when analyzing only the first 5 (or any random five; not shown) trials of each condition. This demonstrates that the CRM task is an effective, robust means with which to induce (and quantify) listening load. There was no correlation between task performance, quantified as the difference in performance between HL and LL conditions, and the PD difference.

### Load-induced vs. Standard measures of pupil responsivity

Figure 4 presents the load-task related pupil responses in Experiment 1 plotted against other standard measures of pupil responsivity. These measures were obtained for each participant to confirm normal pupil reactivity and map out the responsive range, including confirming no ceiling effects. Whilst there was individual variability in pupil size and dilation range, the plot in Figure 4 is a good representation of the average response pattern. In particular, it demonstrates that pupil responses to task-irrelevant sounds (yellow trace) are tiny relative to luminance mediated responses (Pupillary light reflex and pupillary dark reflex in shades of green). Active listening (HL and LL conditions) is associated with increased tonic (baseline) and phasic (evoked by the sentence) pupil dilation. Importantly, the figure also confirms that HL pupil responses are well below ceiling as defined by the maximum pupil dilation measured for each participant.

**Figure 4:**
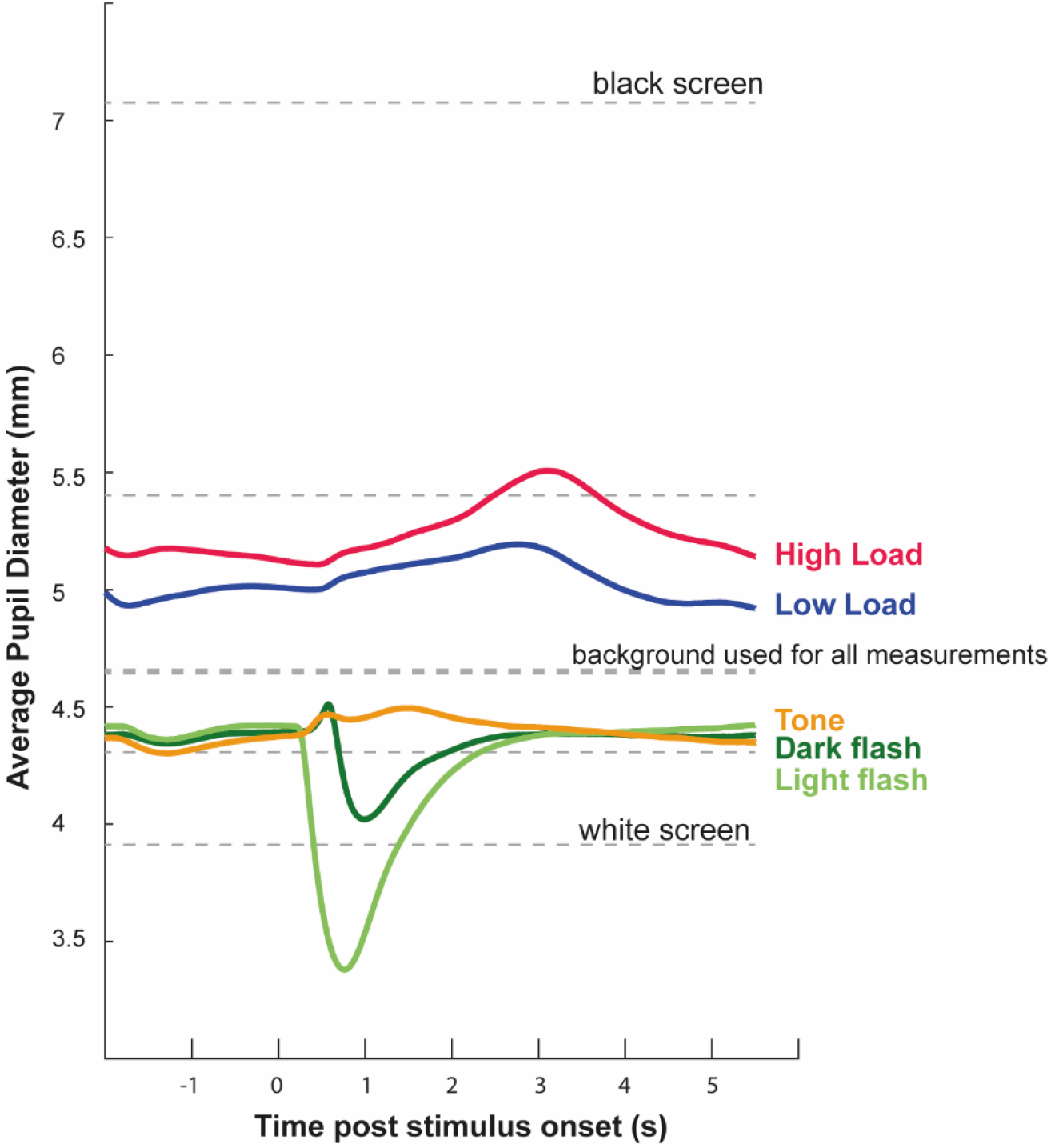
Pupil responsivity. Average (across participants) pupil responsivity in the baseline tasks and the main (speech-in-noise) task in Experiment 1. **Dashed lines**: Participants fixated at the center of the screen while screen brightness changed (10sec intervals; 5 brightness levels) from white to black. Mean pupil diameter in the latter 5 sec of each brightness level are indicated. The screen brightness used as the default background for all other measurements is indicated by the thicker dashed line. **Light green:** Pupillary light reflex. pupil responses to occasional transient (0.3s) screen brightness changes from the default background (mid grey) to white. Average across 30 trials (7 sec inter-flash interval). **Dark green:** Pupillary dark reflex. pupil responses to occasional transient (0.3s) screen brightness changes from the default background (mid grey) to black. Average across 30 trials (7 sec inter-flash interval). **Yellow:** pupil responses to occasional brief (0.3s) harmonic tone. Average across 30 trials (7 sec inter-flash interval). **Blue:** Main task, LL (high SNR) condition. **Red:** Main task, HL (low SNR) condition. The screen brightness test was conducted first; followed by the Tone, Light-, and Dark-reflex tests (in random order).

### Experiment 1: Load-induced microsaccade modulation

Overall, the behavioral and PD data in Experiment 1 established that our task successfully manipulated auditory perceptual load, resulting in conditions that differed in listening effort. Pupil dilation data confirmed that, in line with previous reports, HL conditions were associated with sentence-evoked and sustained effects consistent with increased phasic and tonic arousal under HL. We next turn to the microsaccade rate data.

Figure 5 presents the comparison between the PD data (reproduced from Fig 3) and microsaccade rate data (see Methods), both non-baseline corrected. The MS data exhibit an abrupt microsaccadic inhibition (MSI) response evoked by the onset of the noise, followed by a return to baseline. MSI is commonly observed in response to abrupt sensory events and thought to reflect a rapid interruption of ongoing attentional sampling so as to prioritize the processing of an abrupt sensory event (Roberts et al, 2019; Rolfs et al, 2008; Zhao et al, 2019b). At about 1.5s post-onset, both HL and LL conditions exhibit a drop in MS rate, but this effect is substantially larger for HL: a divergence between the LL and HL conditions is seen between 1.5-2.8s after stimulus onset. Unlike in the PD data where a sustained difference between conditions is seen throughout the epoch, the MS rate effect is confined to a specific period, starting partway through the sentence and ending shortly after target sentence offset.

**Figure 5:**
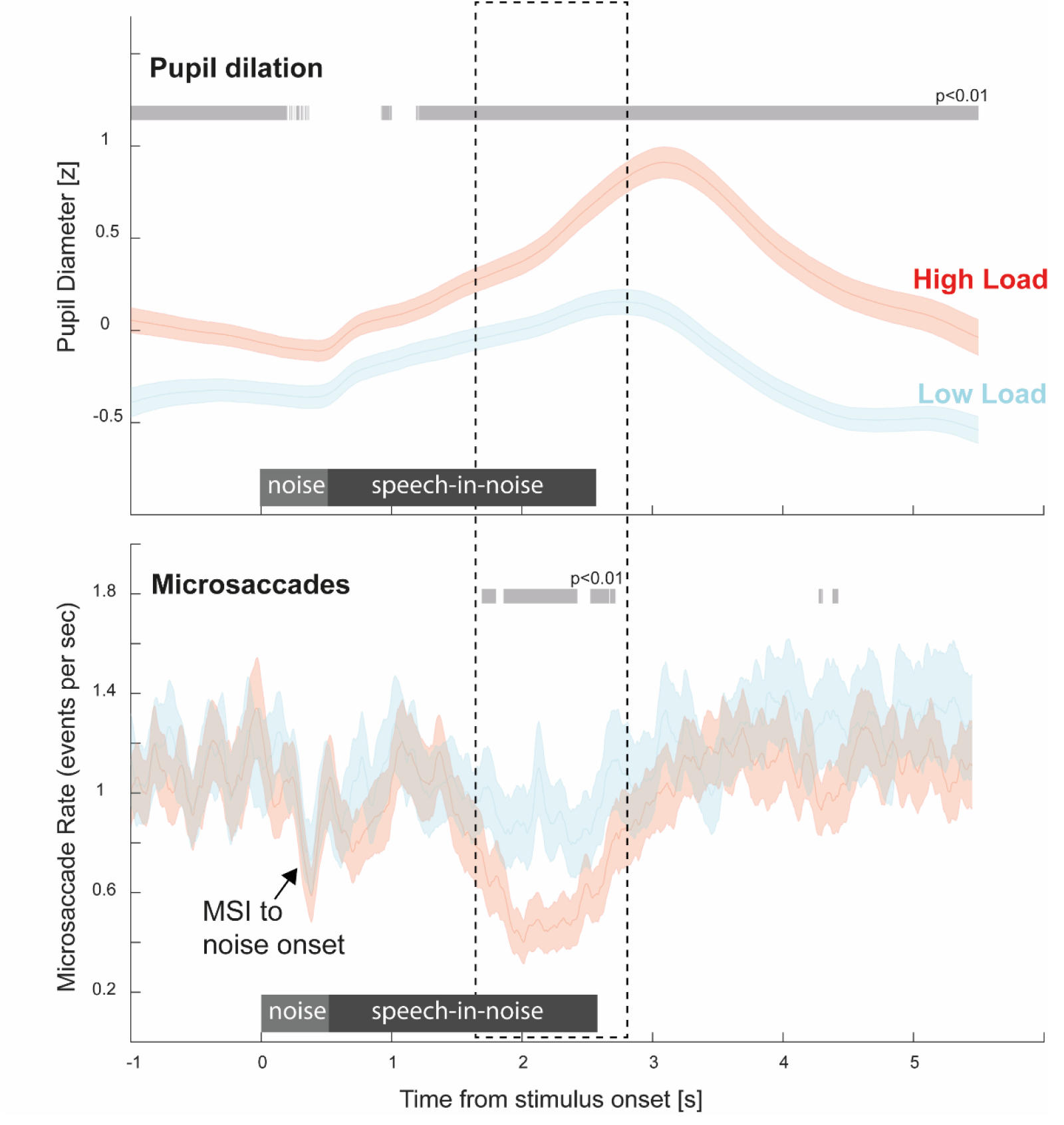
MS incidence is modulated by auditory load. Concurrently recorded PD and MS data. Data are not baseline corrected. Significant differences between conditions are indicated by the grey horizontal lines.

In Experiment 2, we asked whether this effect is linked to the position of the behaviorally relevant elements of the sentence ([color][number] keywords), or rather reflects the time taken for attentional effects to manifest in microsaccadic data. Unlike Experiment 1, where demands on attention (participants focusing to discern the correct color and number) were concentrated at the end of the sentence, ‘Yoda’ sentences in Experiment 2 required focused attention at sentence onset. We hypothesized that if the lower MS rate seen in the HL condition in Experiment 1 reflects a draw on attentional resources, ‘Yoda’ sentences would result in an earlier effect. We also hypothesized that if the timing of the PD peak reflects sentence processing, its latency would similarly be shifted earlier in time in accordance with perceptual demands.

### Experiment 2 – Perceptual load in ‘Yoda’ sentences reveals similar pupil dilation effects to those seen in Experiment 1

‘Yoda’ sentences were created from the original material used in Experiment 1 by moving the keywords ([color], [number]) to the beginning of the sentence (“[color] [number] is, show the dog where the”). To succeed in the task, participants were therefore required to focus attention immediately at sentence onset, but were able to release attention/arousal resources partway through the sentence since later information was not behaviorally relevant.

Figure 6 plots the mean PD to the ‘Yoda’ sentences in noise; z scored (across HL and LL conditions for each participant) and averaged across participants. The pupil dilation responses observed in Experiment 1 were broadly replicated in Experiment 2: A greater PD was observed in the HL condition compared to the LL condition. When baseline correction was applied (Fig 6; left) this effect was significant from ∼1.5s - ∼5s post-onset. In the non-baseline corrected data (Fig 6; right) a sustained difference between conditions was also seen at baseline (from ∼-2 - ∼1.2s pre-onset), though the significant interval was somewhat shorter than that observed in Experiment 1. To elaborate on this, data from Experiment 1 and Experiment 2 were compared directly (Figure 7C).

**Figure 6:**
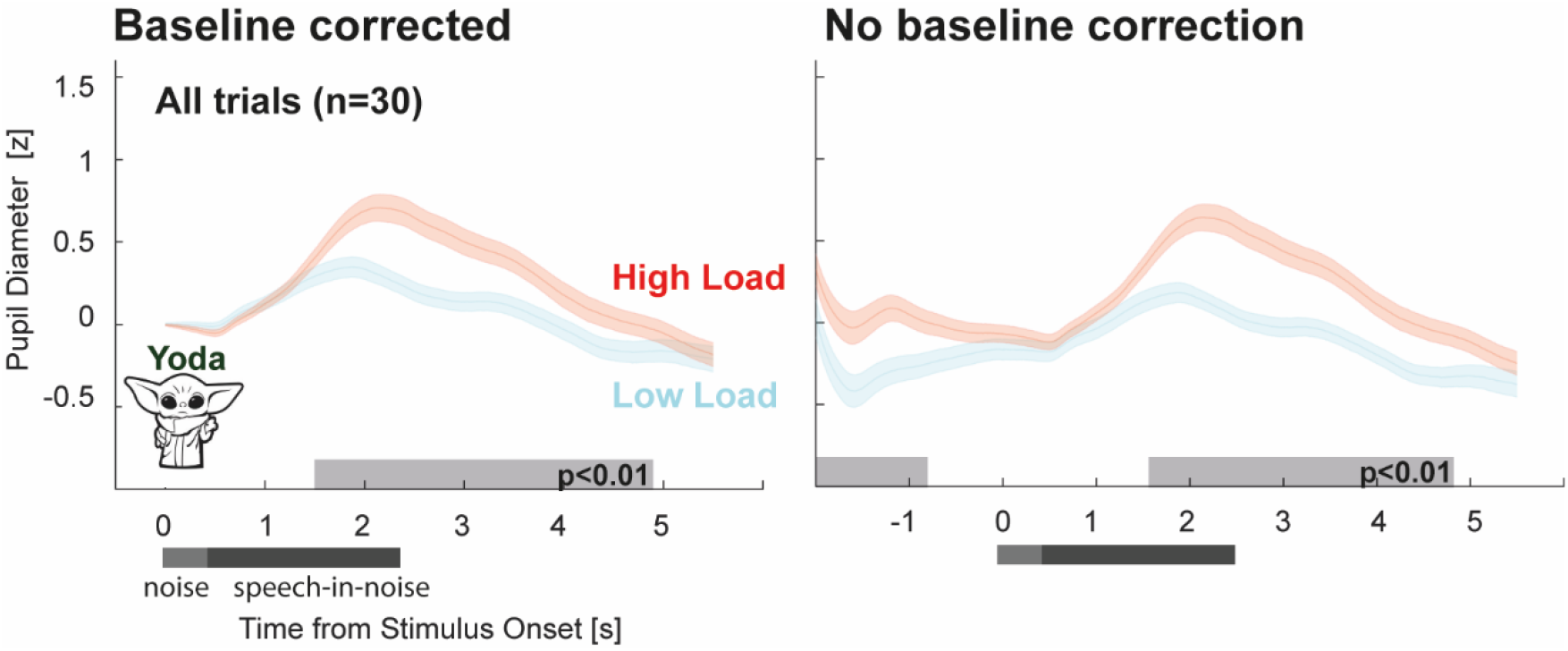
Pupil diameter is modulated by listening load for unusual sentence structures. Pupil diameter was consistently larger in the HL relative to the LL condition. Significant differences between conditions are indicated by the grey horizontal lines. Baseline corrected data on the left; non-baseline corrected on the right. These results replicated the findings of Experiment 1.

### Sentence structure modulates the PD response

Following previous observations that the timing of the PD peak tends to occur after sentence offset, reflecting release of arousal at the conclusion of perceptual processing (Winn et al., 2016; Winn & Moore, 2018; Winn et al., 2022), we hypothesized that the latency of the peak PD in Experiment 2, relative to Experiment 1, would be shifted in accordance with the change in sentence structure. Figure 7 compares the mean PD data across Experiments 1 & 2. A comparison of the HL conditions (non-baseline corrected; Figure 7A) revealed a similar baseline in both experiments. This might be interpreted to suggest a similar effortful state (we return to this point in the discussion section, below). As predicted, Experiment 2 saw the latency of the PD peak shift earlier in time. The comparison between the load conditions (Figure 7C) yielded a similar overall pattern to that observed for HL.

**Figure 7:**
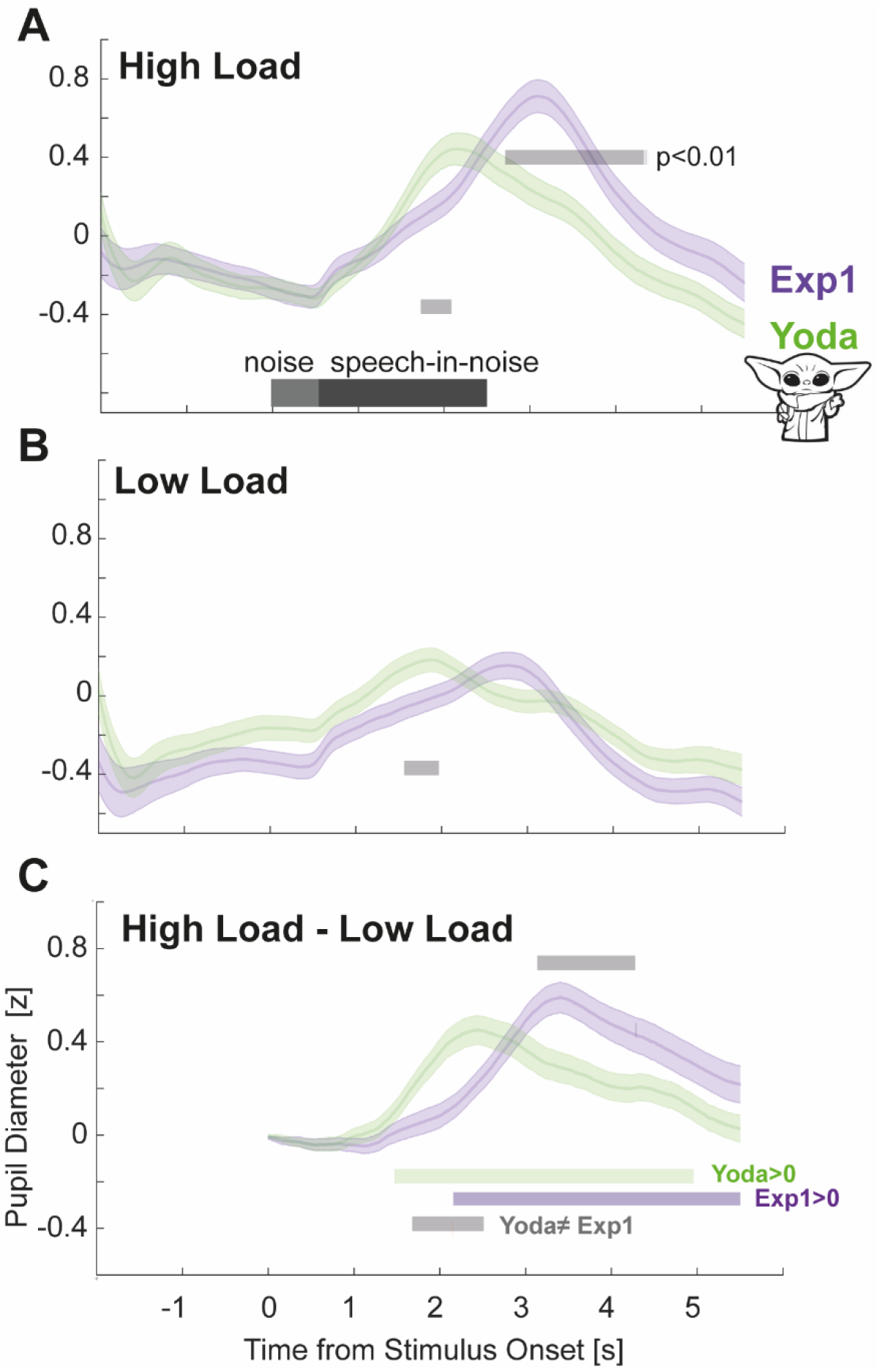
Sentence structure modulates latency of PD response. [A] PD in the HL condition across experiments. Significant differences between experiments are indicated by the grey horizontal lines. [B] PD in the LL condition across experiments. Significant differences between experiments are indicated by the grey horizontal lines. [C] HL-LL Difference between experiments. Green horizontal line: Experiment 2 difference when compared to zero. Purple horizontal line: Experiment 1 difference when compared to zero.

We also observed differences between Experiments 1 and 2 in the peak amplitude of the PD in the HL condition. An independent samples t-test revealed a significant difference in peak amplitude in the HL condition (t(60) = 2.29, p = 0.025) but not in the LL condition (t(60) = -.48, p = 0.63). The pattern of the PD overall suggests a gradual increase in arousal from sentence onset in both experiments with an earlier release (and a lower peak) in Experiment 2.

Figure 7 quantifies the difference between HL and LL in the two experiments by plotting the (baseline corrected) difference waveforms. In Experiment 1, a divergence between HL and LL is seen from ∼2s post onset. In Experiment 2, the two conditions diverge 0.5s earlier (at ∼1.5 s post onset), again consistent with the earlier engagement of arousal in Experiment 2.

### Experiment 2: Load-induced microsaccade modulation

Figure 8 presents the comparison between the PD data (reproduced from Fig 6) and microsaccade rate data, both non-baseline corrected. The MS data exhibit an abrupt microsaccadic inhibition (MSI) response evoked by the onset of the noise. Unlike Experiment 1, this response is not followed by a return to baseline. Microsaccade rate remains low until partway through the sentence in both HL and LL conditions. Thereafter, a return to baseline is observed. This is consistent with the fact that in Experiment 2 attentional resources are required during the initial portion of the sentence. In the LL condition, this inhibition returns to baseline faster than in the HL condition, resulting in a significant difference around ∼1.2-1.5s post-onset. Thus, while a reduction in MS rate was present in both conditions, it lasted longer in the HL condition.

**Figure 8:**
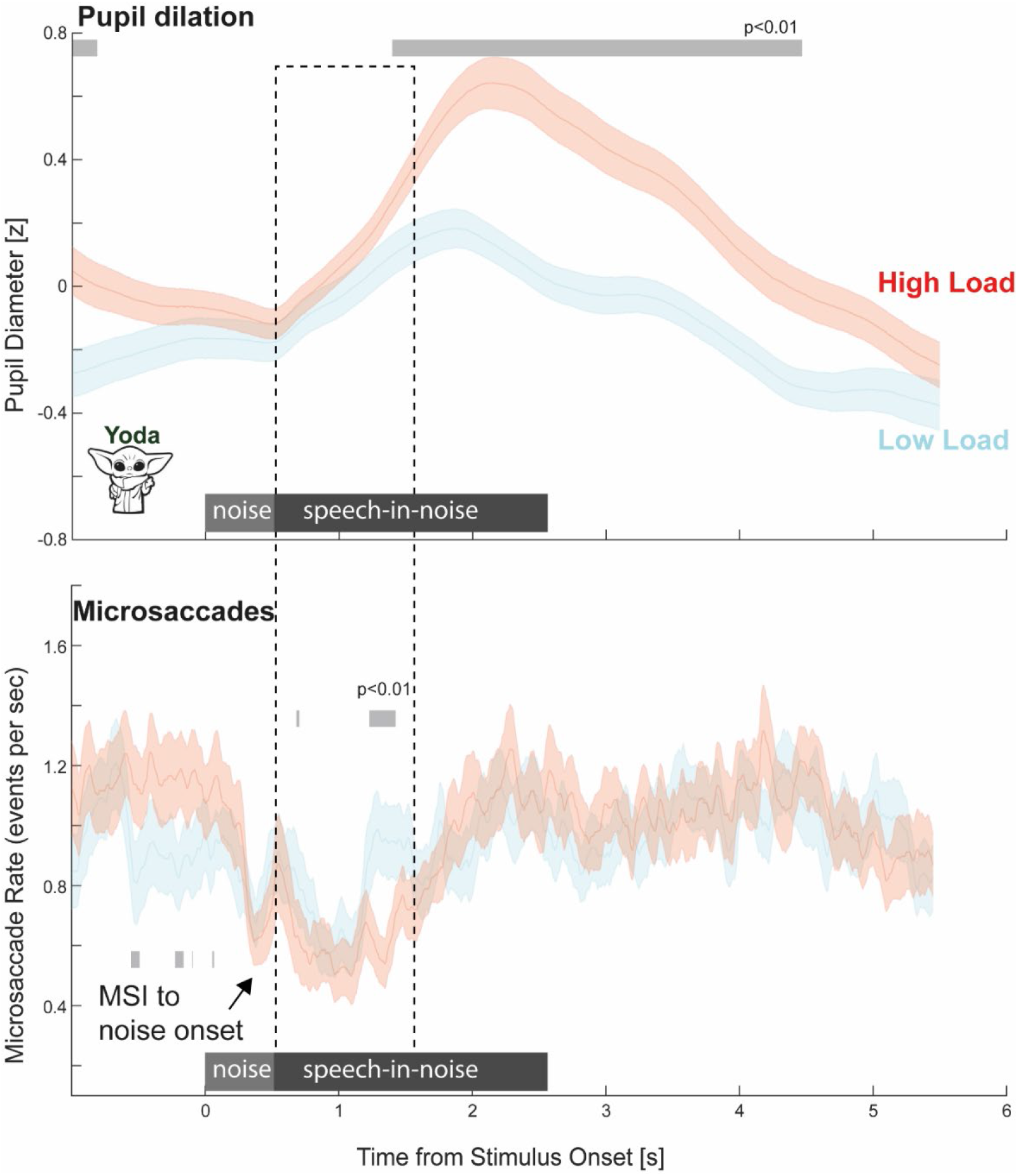
Experiment 2: MS incidence is modulated by auditory load. Concurrently recorded PD and MS data. Data are not baseline corrected. Significant differences between conditions are indicated by the grey horizontal lines

Furthermore, there appears to be an additional effect during the baseline (pre-sentence onset) period where the LL condition exhibits a reduction in MS rate relative to the HL condition. This might reflect the active pre-allocation of attentional resources in this condition. That the effect is only observed in the LL condition is interesting and might imply a depletion of relevant resources in HL that precluded participants from preparing to attend in the same way. We return to this in the discussion section.

### Microsaccade rate modulation reflects localized demand on attention

To directly compare the timing of MS rate modulation, Figure 9 presents data from Experiments 1 & 2, separately for the HL and LL conditions. MS rate was significantly lower in Experiment 2 during the initial portion of the sentence and vice versa during the latter portion of the sentence – consistent with where the keyword coordinates were embedded. This effect is seen in both the HL and LL conditions, but greater (in terms of deflection from baseline) in the HL condition, reflecting the higher demand for attentional resources during low SNR. Critically, in each condition the period of MS modulation (where MS rate was different from baseline) was confined to the period in the sentence where the keywords were present.

**Figure 9:**
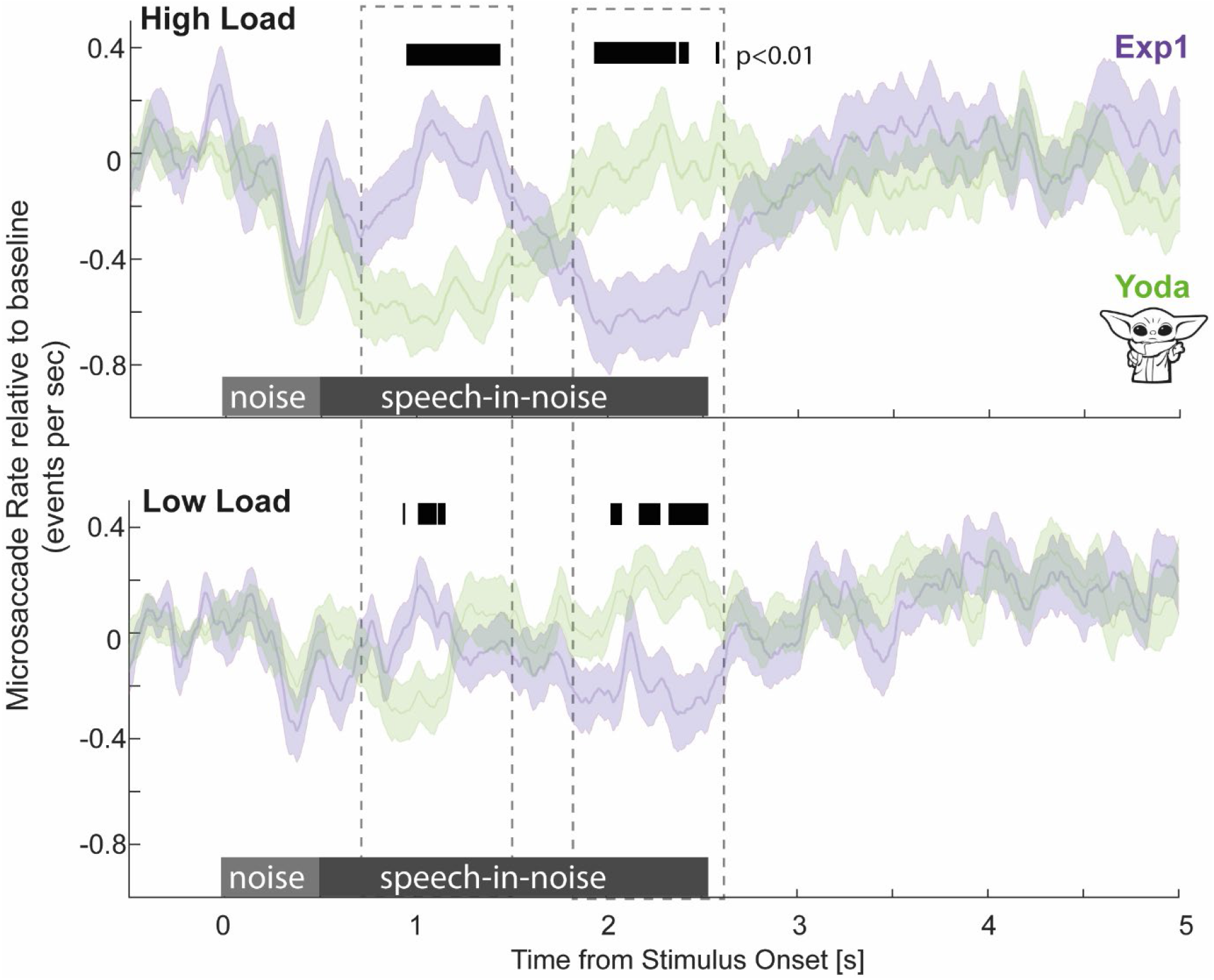
MS rate reflects instantaneous demand on attentional resources. [Top] HL conditions in Experiment 1 and Experiment 2 (“Yoda”). [Bottom] LL conditions in Experiment 1 and Experiment 2 (“Yoda”), all baseline corrected. Significant differences between conditions are indicated with the black horizontal bar.

## Discussion

Successful listening in busy environments requires the deployment of arousal, attention, and other cognitive functions, usually collated under the umbrella-term “listening effort” (Peelle, 2018; Pichora-Fuller, 2016). Quantifying the contribution of each factor to effortful listening is critical for understanding the challenges listeners face in adverse conditions, characterizing individual difficulties (Winn & Teece, 2022) and guiding rehabilitation. We recorded pupil dilation (PD) and microsaccade (MS) rate data while listeners performed a speech-in-noise detection task simulating high- and low-effort listening conditions (HL and LL). PD is a prevalent measure of listening effort (Zekveld et al, 2018; Winn et al, 2018), believed to index modulation of arousal. MS are hypothesized to reflect a process of unconscious visual exploratory sampling, and are therefore a potentially useful signal for understanding how the auditory system interfaces with the brain’s attention network. We hypothesized that measuring MS alongside PD would enable us to precisely identify and measure the deployment of focused attention during speech processing.

Consistent with established literature on PD as an index of listening effort, we observed phasic (sentence evoked) and tonic (sustained) differences in pupil diameter between HL and LL conditions. Unlike the temporally extended PD responses, MS effects, characterized by a reduced MS rate, were only present when demands on focused attention were highest. These results demonstrate that auditory selective attention interfaces with the mechanisms controlling MS generation, establishing MS as an informative measure with which to quantify the temporal dynamics of auditory attentional processing during effortful listening.

### Pupil dilation data reveal tonic and phasic modulation of arousal under load

In line with the established link between PD and listening effort (Zekveld et al, 2018) and consistent with the general association between greater arousal and larger pupil sizes (Bradley et al., 2008; Partala & Surakka, 2003), we observed robust effects of load on the pupillary response. Baseline corrected PD data revealed a significant difference between load conditions, suggesting greater instantaneous arousal under HL. Additional extended differences in non-baseline corrected data are consistent with sustained alertness and engagement under HL.

The latency of the PD peak shifted with the positions of the behaviorally relevant keywords (Fig 7). This is consistent with the notion of ‘effort release’ (Winn et al, 2016; Winn & Moore, 2018; Winn &Teece, 2022; Teece & Winn, 2021). Winn et al (2016; see also McMurray et al., 2017) demonstrated that sentential material of increasing complexity (low semantic context or vocoding) is associated with later decay of PD post-offset, indicative of increased (and presumably more temporally prolonged) demands on processing lasting beyond sentence offset. In Winn & Moore (2018), effort release happened earlier when sentences were followed by ignored stimuli compared to attended digit sequences. Similar effects were observed here with the PD peak occurring earlier for the ‘Yoda’ stimuli (Experiment 2) relative to the original sentences (Experiment 1).

In addition to the shift in latency we also observed that the amplitude of the HL peak in Experiment 1 was larger than that in Experiment 2 (Fig 7A). This reveals that effortful listening is associated with a steady accumulation of arousal as the sentence unfolds. Hence earlier behavioral disengagement (e.g. ‘Yoda’ sentences here) is associated with both an earlier and shallower peak dilation.

In summary, several processes - instantaneous arousal, attentional engagement, and sustained arousal related to the adverse listening environment all appear to contribute to observed PD modulation.

### Microsaccadic activity indexes instantaneous auditory attention

Microsaccades are small spontaneous fixational eye movements occurring at a rate between 1-2 Hz. Accumulating evidence suggests that MS reflect a process of unconscious visual exploratory sampling which modulates early visual processing and plays a critical role in visual perception (Lowet et al, 2018). Importantly, recent findings suggest that MS sampling draws on a central resource pool shared with other perceptual processes, such that MS incidence is affected by the load currently experienced by the individual. MS-indexed visual exploration has been shown to reduce in anticipation of task-relevant events (Denison et al, 2019; Abeles et al, 202), as a function of task engagement (e.g. absorption during music listening; Lange et al, 2017), and under high load (Siegenthaler et al, 2014; Gau et al, 2015; Dalmaso et al, 2017; Yablonski et al, 2017). MS is therefore a useful signal for quantifying participants’ instantaneous attentional state.

Though research into MS dynamics has predominantly focused on visual processing - spatial attention in particular (Engbert, 2006; Krauzlis et al, 2017)- previous demonstrations have linked MS-indexed sampling and auditory processing: anticipation of target sounds causes reduced sustained MS activity (Widman et al, 2014; Abeles et al, 2020) and perceptual salience of brief sounds modulates transient evoked microsaccadic inhibition (Valsecchi & Turatto, 2009; Zhao et al, 2019). Interestingly, abrupt sounds cause more rapid MS inhibition than visual stimuli (Rolfs, 2009), suggesting fast circuitry consistent with the auditory system having privileged access into MS-indexed attention. Here we further demonstrated that instantaneous top-down auditory attention modulates MS dynamics.

Microsaccade activity was recorded concurrently with PD. Since no element of the task was spatialized, we pooled over MS direction and focused on incidence. Unlike the sustained differences between HL and LL conditions observed in the PD data, MS dynamics exhibited a localized effect specifically at points within the sentence containing behaviorally relevant information (keywords). This supports the hypothesis that MS rate reduction reflects an instantaneous draw on attention resources. A similar pattern of MS dynamics was observed in HL and LL conditions (Figure 9) - but the effects relative to baseline are more distinct in HL. This reflects a greater draw on attentional resources exacerbated by the arduous listening environment. Overall, these results demonstrate that MS dynamics can track participants’ instantaneous attentional state: monitoring microsaccade activity during speech processing can reveal how listeners allocate attentional resources to the unfolding sentential material.

In Experiment 2 (Yoda sentences) a smaller difference in MS rate between HL and LL was observed, potentially because the unusual listening demands (keywords exactly at onset) depleted attentional resources in both conditions. Interestingly, whilst MS modulation was largely confined to keyword locations, some pre-onset effects were seen in the LL condition, potentially reflecting preparatory attention allocation to sentence onset. Curiously, this effect was not observed in HL, perhaps because load-induced fatigue depleted listeners’ ability to effectively deploy preparatory attention. This observation could be expanded upon in future research to gain a fine-grained perspective on the dynamics of auditory attention.

### Unravelling arousal and attention effects

The MS and PD effects reported here reflect the coordinated operation of a network that regulates arousal and attention. However, the architecture of this network, and specifically the interrelation between PD and MS control circuits, is the subject of current research.

MS are controlled by a network encompassing the superior colliculus (SC) and frontal eye field (FEF)-an area that plays a key role in controlling attention and distraction (Peele et al, 2016; Lega et al, 2019; Hsu et al, 2021). The SC is considered the dominant driver of MS (Goffart et al., 2012; Hafed, 2011; Hafed & Krauzlis, 2012) and a key structure for visual attention (Herman and Krauzlis, 2017). It receives sensory, cognitive, and arousal inputs from cortical and sub-cortical sources (including from the locus coeruleus (LC), see below) and projects to brainstem premotor circuits to direct the orienting response.

PD is commonly linked to LC activity (Aston-Jones & Cohen, 2005; Breton-Provencher & Sur, 2019; Joshi et al., 2016; Murphy et al., 2014; Reimer et al., 2016) which is at the core of arousal regulation (Aston-Jones & Cohen, 2005; Berridge, 2008; Breton-Provencher & Sur, 2019; Samuels & Szabadi, 2008a, 2008b). The LC has also been implicated in attentional control (Robins, 1984; Aston-Jones et al, 1999) through projections to the pre-frontal cortex (PFC) and SC (Wang et al; 2017; Wang & Munoz 2021; Benavidez et al, 2021). Indeed, LC-NE system responsiveness correlates with enhanced attentional performance (Dahl et al, 2020). Optogenetic activation of LC-NE neurons improves attention and response inhibition via projections to PFC (Bari et al, 2020), with overall attentional state mediated by tonic and phasic LC activity (Aston-Jones & Cohen, 2005; Howells et al, 2012). Therefore, the PD signal likely indexes activity correlating with both attention and arousal.

In line with the multiple links between MS and PD circuits (including via SC and PFC), there are reports of correlations between PD-indexed micro fluctuations of arousal and spontaneous-or SC microstimulation evoked-MS activity (Johnston et al, 2022; Wang & Munoz, 2021; Wang et al, 2022). Similarly, here the modulation of MS rate occurred during the rising slope of PD (where change in pupil size was fastest). This is consistent with attention and arousal interacting to support the perceptual processing of the sentence.

Overall, our results show that whilst the pupil signal is affected by both attention and arousal, the time specificity of MS modulation renders them a powerful tool for pinpointing effects of attention. Future work can capitalize on MS as a critical window into the dynamics of auditory focused attention and its deficits.

## Acknowledgements

CCW and NM are supported by the NIHR UCLH BRC Deafness and Hearing Problems Theme.

## Notes

**Conflict of interest** The authors declare no competing financial interests.

### Competing Interest Statement

The authors have declared no competing interest.

## References

Abeles D, Amit R, Tal-Perry N, Carrasco M, Yuval-Greenberg S (2020) Oculomotor inhibition precedes temporally expected auditory targets. Nat Commun 11:1–12.

Alhanbali S, Dawes P, Millman RE, Munro KJ (2019) Measures of Listening Effort Are Multidimensional. Ear Hear 40:1084–1097.

Aston-Jones G, Rajkowski J, Cohen J (1999) Role of locus coeruleus in attention and behavioral flexibility. Biological Psychiatry 46:1309–1320.

Aston-Jones G, Cohen JD (2005) Adaptive gain and the role of the locus coeruleus– norepinephrine system in optimal performance. Journal of Comparative Neurology 493:99–110.

Bari A, Xu S, Pignatelli M, Takeuchi D, Feng J, Li Y, Tonegawa S (2020) Differential attentional control mechanisms by two distinct noradrenergic coeruleo-frontal cortical pathways. Proceedings of the National Academy of Sciences 117:29080–29089.

Benavidez NL et al. (2021) Organization of the inputs and outputs of the mouse superior colliculus. Nat Commun 12:4004.

Berridge CW (2008) Noradrenergic modulation of arousal. Brain Research Reviews 58:1–17.

Bolia RS, Nelson WT, Ericson MA, Simpson BD (2000) A speech corpus for multitalker communications research. The Journal of the Acoustical Society of America 107:1065–1066.

Bradley MM, Miccoli L, Escrig MA, Lang PJ (2008) The pupil as a measure of emotional arousal and autonomic activation. Psychophysiology 45:602–607.

Breton-Provencher V, Sur M (2019) Active control of arousal by a locus coeruleus GABAergic circuit. Nat Neurosci 22:218–228.

Dahl MJ, Mather M, Sander MC, Werkle-Bergner M (2020) Noradrenergic Responsiveness Supports Selective Attention across the Adult Lifespan. J Neurosci 40:4372–4390.

Dalmaso M, Castelli L, Scatturin P, Galfano G (2017) Working memory load modulates microsaccadic rate. Journal of Vision 17:6.

Dayan P, Abbott LF (2001) Theoretical neuroscience: computational and mathematical modeling of neural systems. Cambridge, Mass: Massachusetts Institute of Technology Press.

Denison RN, Yuval-Greenberg S, Carrasco M (2019) Directing Voluntary Temporal Attention Increases Fixational Stability. J Neurosci 39:353–363.

Efron B, Tibshirani R (1993) An introduction to the bootstrap. New York: Chapman & Hall.

Engbert R, Kliegl R (2003) Microsaccades uncover the orientation of covert attention. Vision Research 43:1035–1045.

Engbert R (2006) Microsaccades: a microcosm for research on oculomotor control, attention, and visual perception. In: Progress in Brain Research (Martinez-Conde s, Macknik SL, Martinez LM, Alonso J-M, Tse PU, eds), pp 177–192 Visual Perception. Elsevier.

Gao X, Yan H, Sun H (2015) Modulation of microsaccade rate by task difficulty revealed through between- and within-trial comparisons. Journal of Vision 15:3.

Goffart L, Hafed ZM, Krauzlis RJ (2012) Visual Fixation as Equilibrium: Evidence from Superior Colliculus Inactivation. Journal of Neuroscience 32:10627–10636.

Hafed ZM (2011) Mechanisms for generating and compensating for the smallest possible saccades. European Journal of Neuroscience 33:2101–2113.

Hafed ZM, Krauzlis RJ (2012) Similarity of superior colliculus involvement in microsaccade and saccade generation. Journal of Neurophysiology 107:1904–1916.

Hafed ZM, Chen C-Y, Tian X (2015) Vision, Perception, and Attention through the Lens of Microsaccades: Mechanisms and Implications. Frontiers in Systems Neuroscience 9

Haro S, Rao HM, Quatieri TF, Smalt CJ (2022) EEG alpha and pupil diameter reflect endogenous auditory attention switching and listening effort. European Journal of Neuroscience 55:1262–1277.

Herman JP, Krauzlis RJ (2017) Color-Change Detection Activity in the Primate Superior Colliculus. eNeuro 4:ENEURO.0046-17.2017.

Hopstaken JF, van der Linden D, Bakker AB, Kompier MAJ (2015) The window of my eyes: Task disengagement and mental fatigue covary with pupil dynamics. Biological Psychology 110:100–106.

Howells FM, Stein DJ, Russell VA (2012) Synergistic tonic and phasic activity of the locus coeruleus norepinephrine (LC-NE) arousal system is required for optimal attentional performance. Metab Brain Dis 27:267–274.

Hsu T-Y, Chen J-T, Tseng P, Wang C-A (2021) Role of the frontal eye field in human microsaccade responses: A TMS study. Biological Psychology 165:108202.

Johnston R, Snyder AC, Khanna SB, Issar D, Smith MA (2022) The eyes reflect an internal cognitive state hidden in the population activity of cortical neurons. Cerebral Cortex 32:3331–3346.

Joshi S, Li Y, Kalwani RM, Gold JI (2016) Relationships between Pupil Diameter and Neuronal Activity in the Locus Coeruleus, Colliculi, and Cingulate Cortex. Neuron 89:221–234.

Krauzlis RJ, Goffart L, Hafed ZM (2017) Neuronal control of fixation and fixational eye movements. Philosophical Transactions of the Royal Society B: Biological Sciences 372:20160205.

LaFrance A (2015) An Unusual Way of Speaking, Yoda Has. The Atlantic Available at: https://www.theatlantic.com/entertainment/archive/2015/12/hmmmmm/420798/ [Accessed February 1, 2023].

Lange EB, Zweck F, Sinn P (2017) Microsaccade-rate indicates absorption by music listening. Consciousness and Cognition 55:59–78.

Lega C, Ferrante O, Marini F, Santandrea E, Cattaneo L, Chelazzi L (2019) Probing the Neural Mechanisms for Distractor Filtering and Their History-Contingent Modulation by Means of TMS. J Neurosci 39:7591–7603.

Lowet E, Gomes B, Srinivasan K, Zhou H, Schafer RJ, Desimone R (2018) Enhanced Neural Processing by Covert Attention only during Microsaccades Directed toward the Attended Stimulus. Neuron 99:207-214.e3.

Messaoud-Galusi Souhila, Hazan V, Rosen S (2011) Investigating Speech Perception in Children With Dyslexia: Is There Evidence of a Consistent Deficit in Individuals? Journal of Speech, Language, and Hearing Research 54:1682–1701.

McGarrigle R, Dawes P, Stewart AJ, Kuchinsky SE, Munro KJ (2017) Pupillometry reveals changes in physiological arousal during a sustained listening task. Psychophysiology 54:193–203.

McGarrigle R, Rakusen L, Mattys S (2021) Effortful listening under the microscope: Examining relations between pupillometric and subjective markers of effort and tiredness from listening. Psychophysiology 58:e13703.

McMurray B, Farris-Trimble A, Rigler H (2017) Waiting for lexical access: Cochlear implants or severely degraded input lead listeners to process speech less incrementally. Cognition 169:147–164.

Murphy PR, O’Connell RG, O’Sullivan M, Robertson IH, Balsters JH (2014) Pupil diameter covaries with BOLD activity in human locus coeruleus. Human Brain Mapping 35:4140–4154.

Ohlenforst B, Wendt D, Kramer SE, Naylor G, Zekveld AA, Lunner T (2018) Impact of SNR, masker type and noise reduction processing on sentence recognition performance and listening effort as indicated by the pupil dilation response. Hearing Research 365:90–99.

Partala T, Surakka V (2003) Pupil size variation as an indication of affective processing. International Journal of Human-Computer Studies 59:185–198.

Peel TR, Hafed ZM, Dash S, Lomber SG, Corneil BD (2016) A Causal Role for the Cortical Frontal Eye Fields in Microsaccade Deployment. PLOS Biology 14:e1002531.

Peelle JE (2018) Listening Effort: How the Cognitive Consequences of Acoustic Challenge Are Reflected in Brain and Behavior. Ear Hear 39:204–214.

Pichora-Fuller MK, Kramer SE, Eckert MA, Edwards B, Hornsby BWY, Humes LE, Lemke U, Lunner T, Matthen M, Mackersie CL, Naylor G, Phillips NA, Richter M, Rudner M, Sommers MS, Tremblay KL, Wingfield A (2016) Hearing Impairment and Cognitive Energy: The Framework for Understanding Effortful Listening (FUEL). Ear and Hearing 37:5S.

Reimer J, McGinley MJ, Liu Y, Rodenkirch C, Wang Q, McCormick DA, Tolias AS (2016) Pupil fluctuations track rapid changes in adrenergic and cholinergic activity in cortex. Nat Commun 7:13289.

Ritz H, Wild CJ, Johnsrude IS (2022) Parametric Cognitive Load Reveals Hidden Costs in the Neural Processing of Perfectly Intelligible Degraded Speech. J Neurosci 42:4619–4628.

Robbins TW (1984) Cortical noradrenaline, attention and arousal1. Psychological Medicine 14:13–21.

Roberts MJ, Lange G, Van Der Veen T, Lowet E, De Weerd P (2019) The Attentional Blink is Related to the Microsaccade Rate Signature. Cerebral Cortex 29:5190–5203.

Rolfs M, Kliegl R, Engbert R (2008) Toward a model of microsaccade generation: The case of microsaccadic inhibition. Journal of Vision 8:5.

Rolfs M (2009) Microsaccades: Small steps on a long way. Vision Research 49:2415–2441.

Rucci M, Victor JD (2015) The unsteady eye: an information-processing stage, not a bug. Trends in Neurosciences 38:195–206.

Samuels ER, Szabadi E (2008) Functional Neuroanatomy of the Noradrenergic Locus Coeruleus: Its Roles in the Regulation of Arousal and Autonomic Function Part I: Principles of Functional Organisation. Current Neuropharmacology 6:235–253.

Samuels ER, Szabadi E (2008) Functional Neuroanatomy of the Noradrenergic Locus Coeruleus: Its Roles in the Regulation of Arousal and Autonomic Function Part II: Physiological and Pharmacological Manipulations and Pathological Alterations of Locus Coeruleus Activity in Humans. Current Neuropharmacology 6:254–285.

Siegenthaler E, Costela FM, McCamy MB, Di Stasi LL, Otero-Millan J, Sonderegger A, Groner R, Macknik S, Martinez-Conde S (2014) Task difficulty in mental arithmetic affects microsaccadic rates and magnitudes. European Journal of Neuroscience 39:287–294.

Valsecchi M, Turatto M (2009) Microsaccadic responses in a bimodal oddball task. Psychological Research 73:23–33.

Wang C-A, Munoz DP (2015) A circuit for pupil orienting responses: implications for cognitive modulation of pupil size. Current Opinion in Neurobiology 33:134–140.

Wang C-A, Blohm G, Huang J, Boehnke SE, Munoz DP (2017) Multisensory integration in orienting behavior: Pupil size, microsaccades, and saccades. Biological Psychology 129:36–44.

Wang C-A, White B, Munoz DP (2022) Pupil-linked Arousal Signals in the Midbrain Superior Colliculus. Journal of Cognitive Neuroscience 34:1340–1354.

White BE, Langdon C (2021) The cortical organization of listening effort: New insight from functional near-infrared spectroscopy. NeuroImage 240:118324.

Widmann A, Engbert R, Schröger E (2014) Microsaccadic Responses Indicate Fast Categorization of Sounds: A Novel Approach to Study Auditory Cognition. J Neurosci 34:11152–11158.

Winn MB (2016) Rapid Release From Listening Effort Resulting From Semantic Context, and Effects of Spectral Degradation and Cochlear Implants. Trends in Hearing 20:2331216516669723.

Winn MB, Moore AN (2018) Pupillometry Reveals That Context Benefit in Speech Perception Can Be Disrupted by Later-Occurring Sounds, Especially in Listeners With Cochlear Implants. Trends in Hearing 22:2331216518808962.

Winn MB, Wendt D, Koelewijn T, Kuchinsky SE (2018) Best Practices and Advice for Using Pupillometry to Measure Listening Effort: An Introduction for Those Who Want to Get Started. Trends in Hearing 22:2331216518800869.

Winn MB, Teece K (2021) The timing of deploying and withholding listening effort, in listeners with normal hearing or with cochlear implants. The Journal of the Acoustical Society of America 150:A340–A340.

Winn MB, Teece KH (2022) Effortful Listening Despite Correct Responses: The Cost of Mental Repair in Sentence Recognition by Listeners With Cochlear Implants. Journal of Speech, Language, and Hearing Research 65:3966–3980.

Yablonski M, Polat U, Bonneh YS, Ben-Shachar M (2017) Microsaccades are sensitive to word structure: A novel approach to study language processing. Sci Rep 7:3999.

Zekveld AA, Heslenfeld DJ, Johnsrude IS, Versfeld NJ, Kramer SE (2014) The eye as a window to the listening brain: Neural correlates of pupil size as a measure of cognitive listening load. NeuroImage 101:76–86.

Zekveld AA, Kramer SE (2014) Cognitive processing load across a wide range of listening conditions: Insights from pupillometry. Psychophysiology 51:277–284.

Zekveld AA, Koelewijn T, Kramer SE (2018) The Pupil Dilation Response to Auditory Stimuli: Current State of Knowledge. Trends in Hearing 22:2331216518777174.

Zhao S, Yum NW, Benjamin L, Benhamou E, Yoneya M, Furukawa S, Dick F, Slaney M, Chait M (2019) Rapid Ocular Responses Are Modulated by Bottom-up-Driven Auditory Salience. J Neurosci 39:7703–7714.

Zhao S, Chait M, Dick F, Dayan P, Furukawa S, Liao H-I (2019) Pupil-linked phasic arousal evoked by violation but not emergence of regularity within rapid sound sequences. Nat Commun 10:4030.

